## Supplementary Figures for "Capsules and their traits shape phage susceptibility and plasmid conjugation efficiency"

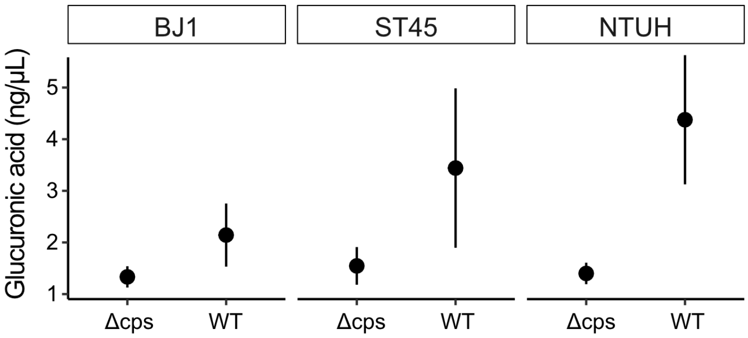


**Figure S1 – Capsule quantification.** Glucuronic acid concentration in capsule extracts (See *Glucuronic acid assay*) in non-capsulated mutants (Δcps) and in the corresponding WT strains. The points represent the average of three independent replicates and the error bar correspond to the standard deviation.

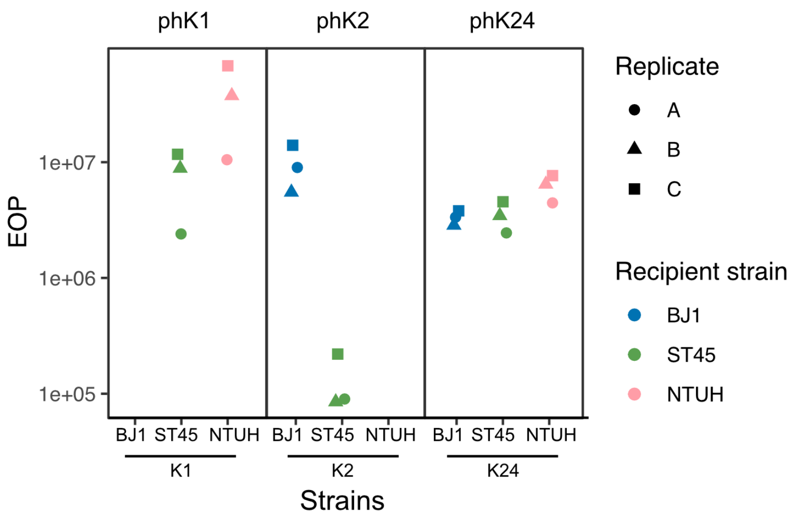


**Figure S2 – Phage infection assay.** Raw values of efficiency of plating (EOP) for each swapped strain. Replicates A, B and C correspond to independent phage lysate prepared with the wildtype strains. Combinations not shown do not result in productive infections (*e.g.* phK1 against ST45::K2).

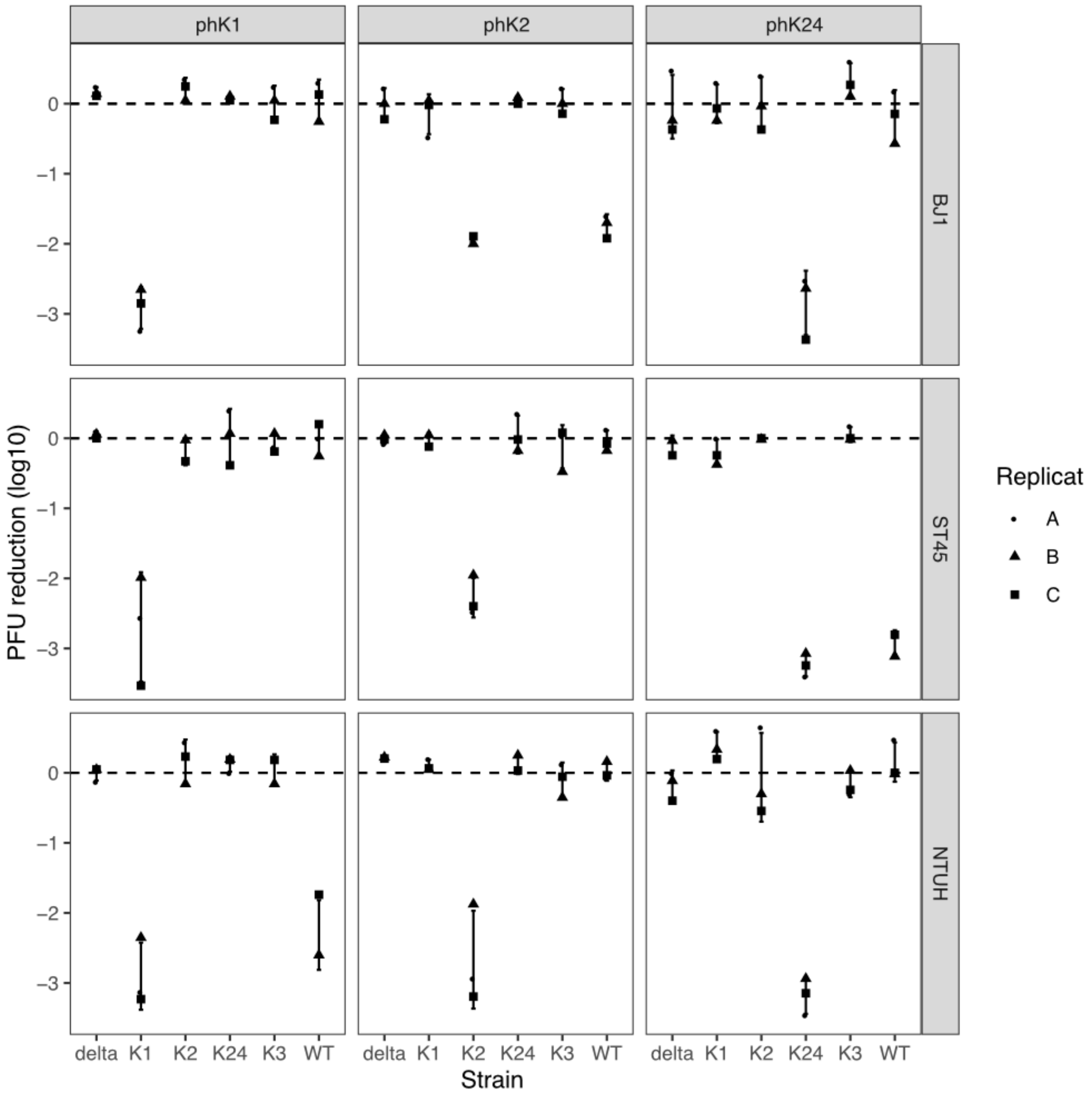


**Figure S3 – Adsorption assay.** Phage adsorption onto the different strains. Shapes correspond to independent replicates. Adsorption is quantified as the log-transformed relative PFU reduction after 5 minutes. A value of 0 indicates no adsorption.

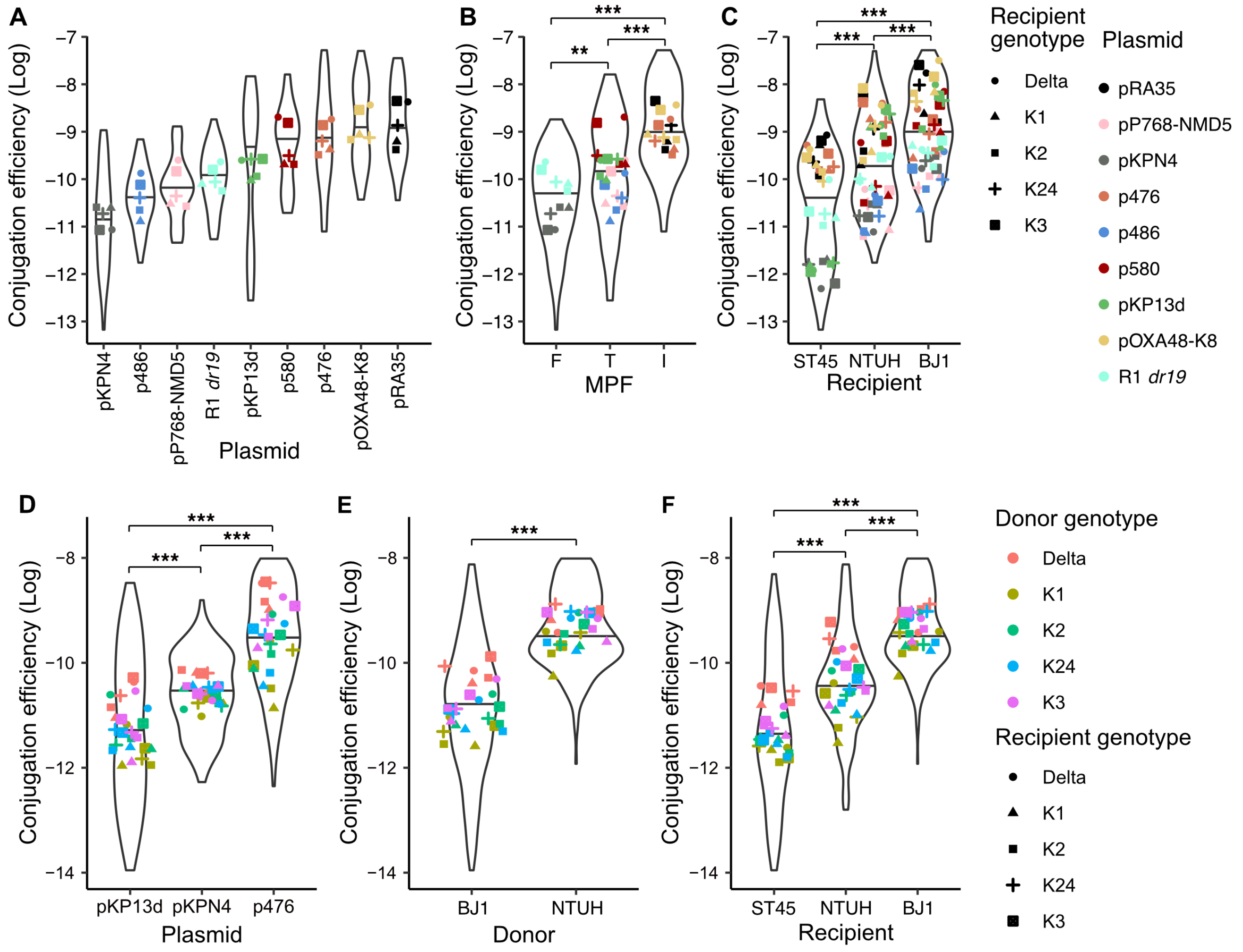


**Figure S4 – Factors accounted for in the multi-variate analysis.**

**A, B, C.** *E. coli* to *K. pneumoniae* assays (set E1).

**D, E, F.** *K. pneumoniae* to *K. pneumoniae* assays (set E2).

A. Log-transformed conjugation efficiency of each plasmid. Individual points are coloured according to the plasmid and represent the average of each recipient genotype (shape).
B. Same data as in A, according to the different MPF types of the plasmids (x-axis).
C. Log-transformed conjugation efficiency of each plasmid (colour) according to the recipient genotype (shape) and recipient strain (x-axis).
D. Log-transformed conjugation efficiency for each plasmid (x-axis) according to the donor (colour) and recipient (shape) genotype.
E. Same data, according to the different donor strains (x-axis).
F. Same data, according to the different recipient strains (x-axis).

Statistical tests: Pairwise Wilcoxon tests. *** $p$<0.001; ** $p$<0.01

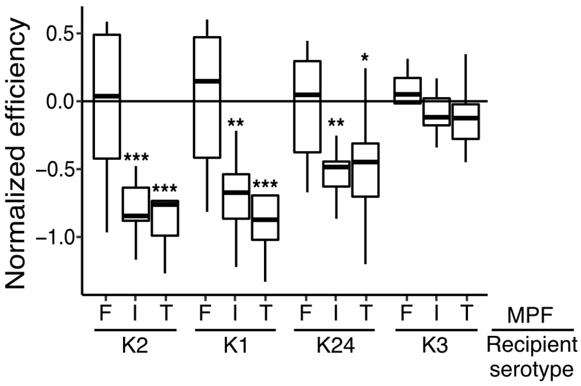


**Figure S5 – Conjugation efficiency of MPF types in distinct recipient serotype**

Drawn from the *E. coli* to *K. pneumoniae* assays (Set E1). The normalized efficiency corresponds to the log-transformed conjugation efficiency of each plasmid-recipient pair subtracted with the one of the corresponding Δcps mutant:

$$Normalized\ efficiency_{Plasmid-Recipient} = \log(Conj.efficiency_{\mathrm{cps+}}) - \log(Conj.efficiency_{\Delta\mathrm{cps}})$$

Below the solid line at 0, the capsulated cells have a lower conjugation efficiency than the Δcps. At 0, there is no difference with the Δcps. Above 0, the capsulated cells have higher conjugation efficiency than the Δcps.

Statistical tests: Wilcoxon test (H0: average =0). *** $p$<0.001; ** $p$<0.01; * $p$<0.05

**Figure S6 – Alignment of OmpK36 proteins from the three *K. pneumoniae* strains.** Proteins aligned with Clustal Omega (https://www.ebi.ac.uk/Tools/msa/clustalo/).

```
OmpK36-BJ1       MKVKVLSLLVPALLVAGAANAAEIYNKDGNKLDLYGKIDGLHYFSDDKSVDGDQTYMRVG     60
OmpK36-NTUH      MKVKVLSLLVPALLVAGAANAAEIYNKDGNKLDLYGKIDGLHYFSDDKSVDGDQTYMRVG     60
OmpK36-ST45      MKVKVLSLLVPALLVAGAANAAEIYNKDGNKLDLYGKIDGLHYFSDDKSVDGDQTYMRVG     60
                 ************************************************************

OmpK36-BJ1       VKGETQINDQLTGYGQWEYNVQANNTESSSDQAWTRLAFAGLKFGDAGSFDYGRNYGVVY     120
OmpK36-NTUH      VKGETQINDQLTGYGQWEYNVQANNTESSSDQAWTRLAFAGLKFGDAGSFDYGRNYGVVY     120
OmpK36-ST45      VKGETQINDQLTGYGQWEYNVQANNTESSSDQAWTRLAFAGLKFGDAGSFDYGRNYGVVY     120
                 ************************************************************

                             L3 LOOP
OmpK36-BJ1       DVTSWTDVLPEFGGDTYGSDNFLQSRANGVATYRNSDFFGLVDGLNFALQYQGKNGSVSG     180
OmpK36-NTUH      DVTSWTDVLPEFGGDTYGSDNFLQSRANGVATYRNSDFFGLVDGLNFALQYQGKNGSVSG     180
OmpK36-ST45      DVTSWTDVLPEFGGDTYGSDNFLQSRANGVATYRNSDFFGLVDGLNFALQYQGKNGSPSG     180
                 ********************************************************* **

                     L4 LOOP
OmpK36-BJ1       EGA---TNNGRGWSKQNGDGFGTSLTYDIWDGISAGFAYSHSKRTDEQNSVPALGRGDNA     237
OmpK36-NTUH      EGA---TNNGRGWSKQNGDGFGTSLTYDIWDGISAGFAYSHSKRTDEQNSVPALGRGDNA     237
OmpK36-ST45      EGALSPTNNGRTALKQNGDGYGTSLTYDIYDGISAGFAYSNSKRLGDQNSKLALGRGDNA     240
                 ***   *****   ******:********:**********:*** .:***  ********

OmpK36-BJ1       ETYTGGLKYDANNIYLASQYTQTYNATRAGSLGFANKAQNFEVVAQYQFDFGLRPSVAYL     297
OmpK36-NTUH      ETYTGGLKYDANNIYLASQYTQTYNATRAGSLGFANKAQNFEVVAQYQFDFGLRPSVAYL     297
OmpK36-ST45      ETYTGGLKYDANNIYLATQYTQTYNATRAGSLGFANKAQNFEVVAQYQFDFGLRPSVAYL     300
                 *****************:******************************************

OmpK36-BJ1       QSKGKDLERGYGDQDILKYVDVGATYYFNKNMSTYVDYKINLLDDNSFTRNAGISTDDVV     357
OmpK36-NTUH      QSKGKDLERGYGDQDILKYVDVGATYYFNKNMSTYVDYKINLLDDNSFTRNAGISTDDVV     357
OmpK36-ST45      QSKGKDLE-GYGDQDILKYVDVGATYYFNKNMSTYVDYKINLLDDNSFTHNAGISTDDVV     359
                 ******** ****************************************:**********

OmpK36-BJ1       ALGLVYQF 365
OmpK36-NTUH      ALGLVYQF 365
OmpK36-ST45      ALGLVYQF 367
                 ********
```
